# Persistent selection on size explains micro- and macroevolutionary alignments in fly wings

**DOI:** 10.64898/2026.02.19.706930

**Authors:** Haoran Cai

## Abstract

The alignment among mutational variance (*M*), standing genetic variance (*G*), and macroevolutionary divergence (*R*) in *Drosophila* wing shape poses a rate paradox under a simple constraint hypothesis: evolution follows mutational lines of least resistance, yet proceeds orders of magnitude slower than the abundant genetic variation would permit. This is difficult to reconcile with a simple constraint view in which long-term evolution merely tracks the amount of available variation in each direction. Previous explanations invoke deleterious pleiotropy on unmeasured traits or correlational selection on trait combinations, but recent empirical work finds little evidence of fitness costs beyond flight performance. Here, by reanalyzing published data, I show that wing size shows the hallmark of the primary selection target: among all wing traits, size exhibits the lowest ratio of standing genetic to mutational variance, indicating the strongest selective depletion. Based on this empirical observation, I develop a single-axis selection model in which natural selection targets only a single trait while all other traits evolve as correlated byproducts via within-module pleiotropy. This minimal model reproduces both the observed *M* –*G*–*R* alignment and slower-than-neutral divergence rates, explaining micro- and macroevolutionary patterns in fly wings without invoking complex adaptive landscapes.

**Significance Statement:** A fundamental question in biology is whether large-scale evolutionary patterns arise from the same processes that drive change within populations. Fly wing shape offers a striking test case: the directions of genetic variation and species divergence are closely aligned, yet wing shape evolves far more slowly than expected — a rate paradox under a simple constraint hypothesis. I show that this paradox can be explained by a simple mechanism: natural selection on one trait indirectly constrains the evolution of other non-selected traits through within-module pleiotropy. This single-axis selection model reproduces observed evolutionary patterns without requiring complex multivariate selection, and its prediction that wing size is the primary target of selection is supported by reanalysis of published data. The framework applies broadly to any trait that shares mutations with correlated characters.

## Introduction

A persistent question in evolutionary biology concerns the relationship between microevolution and macroevolution: can macroevolutionary patterns be explained by microevolutionary processes, and conversely, can population-level changes be predicted from macroevolutionary trends ^1–7^? Microevolution refers to the genetic and phenotypic changes within populations driven by mutation, selection, migration, and drift. Macroevolution describes the broader dynamics of life, including the origination and extinction of lineages and large-scale patterns of phenotypic divergence. While attempts to quantitatively link these two scales have yielded mixed success, a comprehensive framework explaining these variable outcomes remains elusive ^8^.

A notable success in aligning these scales comes from the evolution of wing shape in Drosophila and other Diptera. In this system, a strong positive correlation is consistently observed between the structure of mutational variance (the M-matrix), standing additive genetic variance (the G-matrix), and rates of macroevolutionary divergence (the R-matrix) ^9^. This alignment across different evolutionary timescales offers compelling evidence that non-random phenotypic variation generated by mutation is profoundly linked to long-term evolutionary outcomes.

This alignment, however, presents a paradox under a simple constraint view: if evolutionary trajectories simply follow lines of least resistance, they should proceed much faster given the abundant genetic variation available. Instead, observed macroevolutionary rates are orders of magnitude slower than theoretically permitted, which challenges the simple constraint-based hypothesis ^9^.

One proposed resolution is that most mutational variation is effectively unusable due to deleterious pleiotropic effects on unmeasured traits ^9^. This ‘pleiotropic constraint’ hypothesis predicts genetic covariation between wing shape and fitness components that are functionally unrelated to wing shape or flight. However, an indirect evaluation of this hypothesis in *S. punctum* found little evidence for such covariances, although the authors of that study note their power to detect subtle associations was limited ^10^.

Notably, the slow observed rates challenge the simple constraint explanation but pose no difficulty for selection-based accounts: if selection actively channels evolution along particular directions, slow rates are an expected consequence rather than a puzzle. A second, equally critical aspect is that the alignment between divergence and variation persists over timescales far longer than expected if the direction of selection were arbitrary with respect to *M* and *G*. Several alternative hypotheses have accordingly been proposed:

1. **Moving corridor model** (also known as ‘selective-lines-of-least-resistance’): Net directional selection is concentrated along a major phenotypic axis, while stabilizing selection limits departures in orthogonal directions. The multivariate fitness optimum wanders over geological time, but its trajectory is constrained to “corridors” that roughly parallel the primary axis of mutational variance ^11,12^.
2. **Correlational selection**: Persistent selection on trait combinations (e.g., allometric relationships) shapes both the mutational and genetic variance-covariance matrices, so that divergence naturally aligns with the direction of covariation maintained by selection.
3. **Multidimensional fluctuating selection** (MFS): Under MFS, the multivariate optimum fluctuates rapidly in multiple directions^3^. Because the tracking response is mediated by the *G* matrix, populations take larger evolutionary excursions along axes of high genetic variance. When these rapid fluctuations are time-averaged, the resulting divergence matrix (*R*) naturally aligns with *G* and *M* .

Here, I propose a single-axis selection model as a parsimonious alternative to these hypotheses in fly wing evolution. Like the above models, this model assumes selection is invoked, but posits that selection is concentrated primarily on a single axis. This framework asks whether complex adaptive lanscape or multivariate selection is necessary to reproduce the observed patterns. Using individual-based simulations parameterized with empirical mutational variance-covariance matrices, I show that the single-axis selection model is sufficient to reproduce the observed *M* -*G*-*R* alignment and generate slower-than-neutral divergence rates. Furthermore, empirical evidence suggests that wing size is the primary target of selection, effectively experiencing the strongest selection pressure among all wing traits.

## Results

### Wing Size as the Primary Selection Target

Wing size is a strong candidate for the primary selection target among wing traits, because it directly determines aerodynamic performance: lift scales with wing surface area, so changes in wing size alter flight efficiency. To test this expectation, I used the empirical *G* and *M* matrices estimated for *D. melanogaster* wing traits from ref. ^9^ and computed the per-trait ratio *G*_*ii*_*/M*_*ii*_, which measures how much standing genetic variance is maintained per unit of mutational input. Under pure drift, this ratio should be approximately equal for all traits; persistent selection on a trait preferentially erodes its genetic variance and depresses its *G/M* ratio below the neutral baseline. The *G/M* ratio for wing size is the lowest of all 25 traits, under both the homozygous (*G/M*_hom_ = 72; shape median = 347) and heterozygous (*G/M*_het_ = 144; shape median = 842) mutational variance estimates (Fig. 1). Size retains only *≥*21% (*M*_hom_) to *≥*17% (*M*_het_) as much standing genetic variance per unit of mutational input as the median shape trait. Because standing genetic variance reflects a balance between mutational input and selective removal, the uniquely low *G/M* ratio of size is consistent with it being the primary target of selection, effectively under the strongest stabilizing selection.

**Figure 1:**
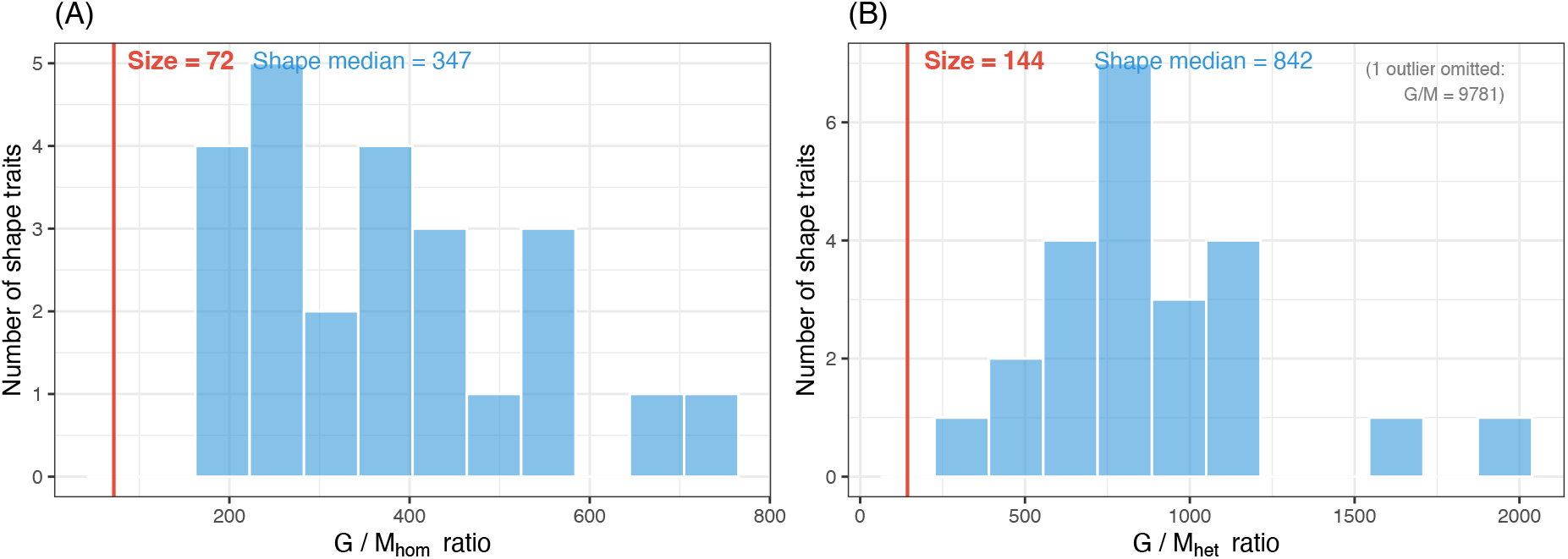
Empirical *G/M* ratio for size vs. shape traits in *D. melanogaster*. Distribution of per-trait *G*_*ii*_*/M*_*ii*_ ratios across the 24 shape traits (blue histogram) computed from the empirical *G* and *M* matrices of ref. ^9^. The vertical red line marks the *G/M* ratio of centroid size (ln CS). (A) *G/M*_hom_ (homozygous mutational variance): size ratio = 72, shape median= 347; (B) *G/M*_het_ (heterozygous mutational variance): size ratio = 144, shape median = 842. Under both estimates, size has the lowest *G/M* ratio of all 25 traits (rank 1/25), consistent with stronger stabilizing selection on size selectively depleting its standing genetic variance.

### Single-Axis Selection Recapitulates the Empirical Alignment Between Divergence and Variation

Based on this observation, here, I propose a single-axis selection model in which only one trait is under persistent selection and all others evolve as correlated byproducts of within-module pleiotropy. Given stable *M* over macroevolutionary time, the divergence matrix *R* should align with both *M* and *G*.

To test whether such a single-axis selection model can reproduce the empirical alignment patterns observed in fly wings, I conducted individual-based simulations using the empirical mutational variance-covariance matrix *M* from ref. ^9^. The simulation protocol was as follows: 20 replicate populations were initialized from an ancestral population experiencing a burn-in session. To allow populations to diverge, each replicate was assigned an optimum moving rate drawn from a normal distribution with mean zero and standard deviation *σ*. This means the optimum translates at a constant velocity for a given lineage, *θ* _*i*_(*t*) = *v*_*i*_*t*, effectively generating a sustained displacement from the optimum rather than a Brownian motion random walk. This design enables populations to track different selective targets at different speeds, generating among-lineage divergence in the primary trait (e.g., size). Larger *σ* values produce faster and more variable divergence among replicates, while smaller *σ* values constrain populations to remain closer to the ancestral optimum. The divergence matrix *R* was computed from the among-population variance in trait means, while the within-population genetic variance-covariance matrix *G* was recorded at the final generation and averaged across 20 replicate populations. Wing size was designated as the primary trait under persistent selection with a moving optimum.

To quantify the alignment between matrices, I employed common subspace analysis following previous approaches ^13,14^. This method projects the matrices onto a common set of orthogonal axes (e.g., the eigenvectors of *M*) and compares the variance along each axis on a logarithmic scale. A strong linear relationship indicates proportional scaling of variances—the signature of alignment between matrices. The following alignment analyses exclude wing size from the matrices, so that the common subspace comparisons specifically test whether selection on size induces proportional scaling of the multivariate covariance structure among the 24 indirectly-selected shape dimensions (including size in the matrices retains similarly strong alignment).

The results show that the single-axis model selecting on size successfully reproduces the empirical patterns (Fig. 2). When comparing log_10_-transformed variances along the principal axes of *M*, both *R* and *G* showed strong positive correlations with *M* . This alignment was robust across two orders of magnitude variation in selection strength on the primary trait (Fig. S6) and across different reference matrices (Fig. S7). Notably, such alignment as the model prediction does not rely on selecting on the wing size but is robust when selecting on wing shape (See Appendix C, Fig. S8). These findings indicate that a model of univariate selection, combined with stable mutational covariances, is sufficient to generate the observed *M* -*G*-*R* alignment for fly wing without invoking complex multivariate selection or correlational selection on shape.

**Figure 2:**
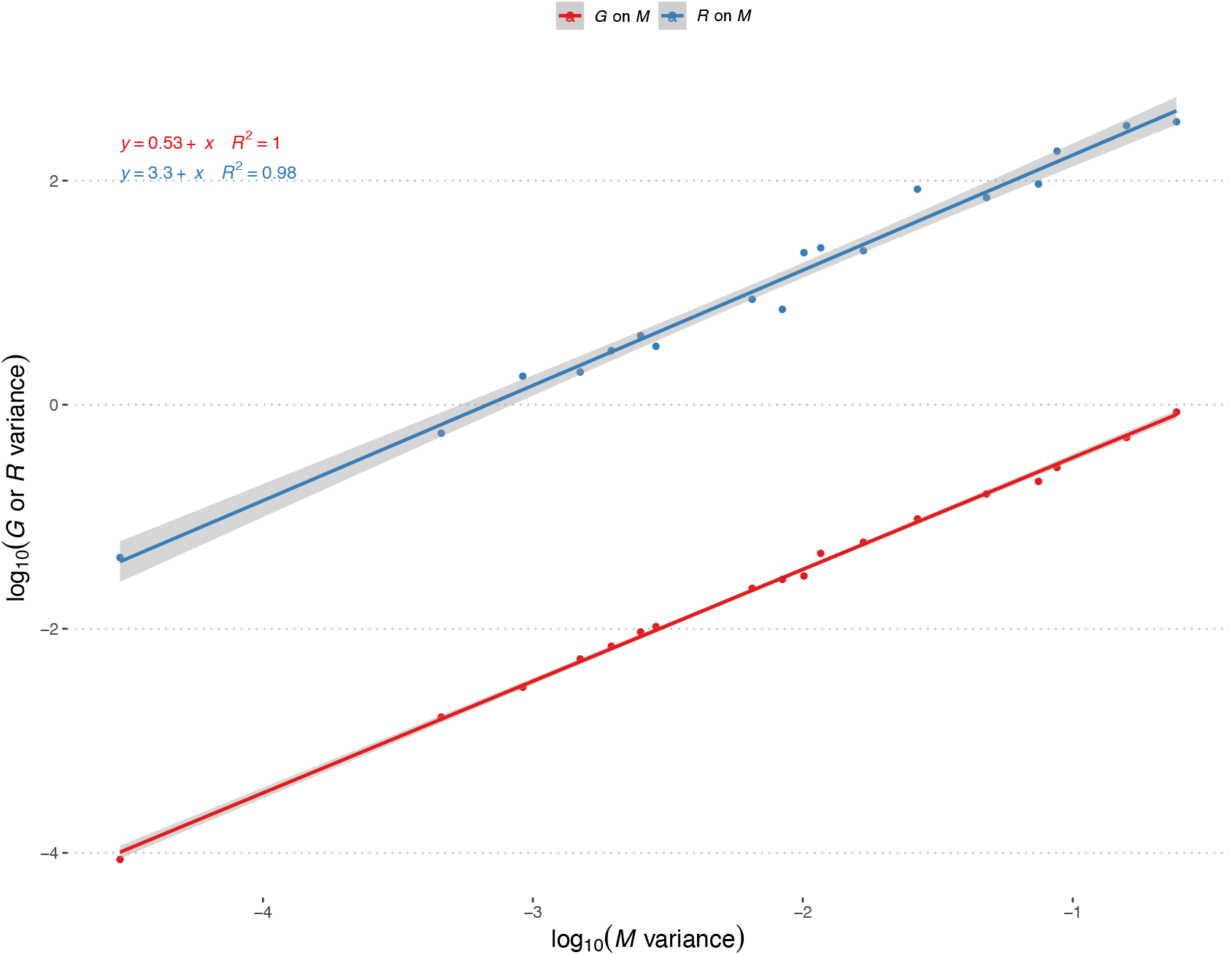
The single-axis selection model reproduces the empirical alignment between *M, G*, and *R*. Common subspace analysis comparing the divergence matrix *R* (blue) and genetic variance matrix *G* (red) to the mutational variance matrix *M* . Points represent log_10_ variance along the eigenvectors of *M* ; lines indicate ordinary least squares regression. Strong positive relationships (*R*^2^ > 0.8) indicate that both *R* and *G* are aligned with the mutational architecture. Only the upper 18 eigenvectors of *M* were used; wing size was not included in the analyses. Shaded areas show 95% CI. Simulations used *V*_*s*_ = 0.05 on wing size (strong selection), *σ* = 0.0001, *N* = 500, and 20 replicate populations evolved for 20,000 generations.

### Depletion of Genetic Variance in the Selection Target

To confirm that the targeted depletion of genetic variance in size from empirical observation (Fig. 1) is a direct expectation of the single-axis selection model, I examined the ratio *G*_*ii*_*/M*_*ii*_ for each trait *i* in the simulated populations. This ratio reflects the amount of standing genetic variance maintained per unit of mutational input.

Results from individual-based simulations support this expectation from the model (Fig. 5). Under neutral evolution, the *G/M* ratio is approximately uniform across all traits, including size, and hovers around ∼ 1,000— closely matching the theoretical expectation of *G/M* = 2*N*_*e*_ = 2 *θ* 500 = 1,000 under pure drift. Under the single-axis selection model in which selection targets size, the size’s *G/M* ratio is markedly reduced compared to shape traits, consistent with selection selectively eroding genetic variance along the selected axis. Furthermore, this *G/M* ratio profile is diagnostic of which trait is under direct selection: when the simulation is repeated with selection on a shape trait instead of size, the lowest *G/M* ratio shifts to that shape trait (Appendix C, Fig. S9).

### Reduced Rate of Shape Divergence Under Single-Axis Selection

The alignment between *M, G*, and *R* alone does not distinguish single-axis selection from completely neutral evolution: under complete neutrality, divergence would also scale with mutational variance ^9,14^. The critical observation that requires explanation is that wing shape evolves orders of magnitude more slowly than expected from the abundant genetic variation available.

The single-axis selection model offers a resolution to this rate paradox. Although shape traits are not directly selected, they experience apparent stabilizing selection through their mutational covariance with wing size. The net result is a strong constraint on shape divergence despite the absence of direct selection. I use the term “divergence rate” to describe the rate of increase in among-lineage variance for these shape traits due to indirect selection responses, mutation, and drift.

Using a simplified two-trait model, I show that selection on the primary trait constrains the drift of correlated neutral traits by reducing the genetic variance of the neutral trait (*G*_*BB*_), thereby reducing the total variance available for drift (see Appendix A for details). Simulation results using the full empirical *M* matrix support these predictions. Figure 3 shows the divergence rate for all non-selected wing shape traits under single-axis selection. Under stronger stabilizing selection on wing size (*V*_*s*_ = 0.05), the divergence rate of every shape trait is substantially reduced (by approximately two orders of magnitude) compared to both weak selection (*V*_*s*_ = 0.5) and the completely neutral scenario, indicating that selection on a single primary trait can constrain the drift of correlated neutral traits. Analogous patterns emerge in the two-trait toy model, where varying selection strength systematically modulates the effective divergence rate (Fig. S3). Moreover, the relationship between mutational variance and divergence rate across traits (Fig. 4) shows that traits with higher mutational input diverge faster, preserving the rate–variance alignment.

**Figure 3:**
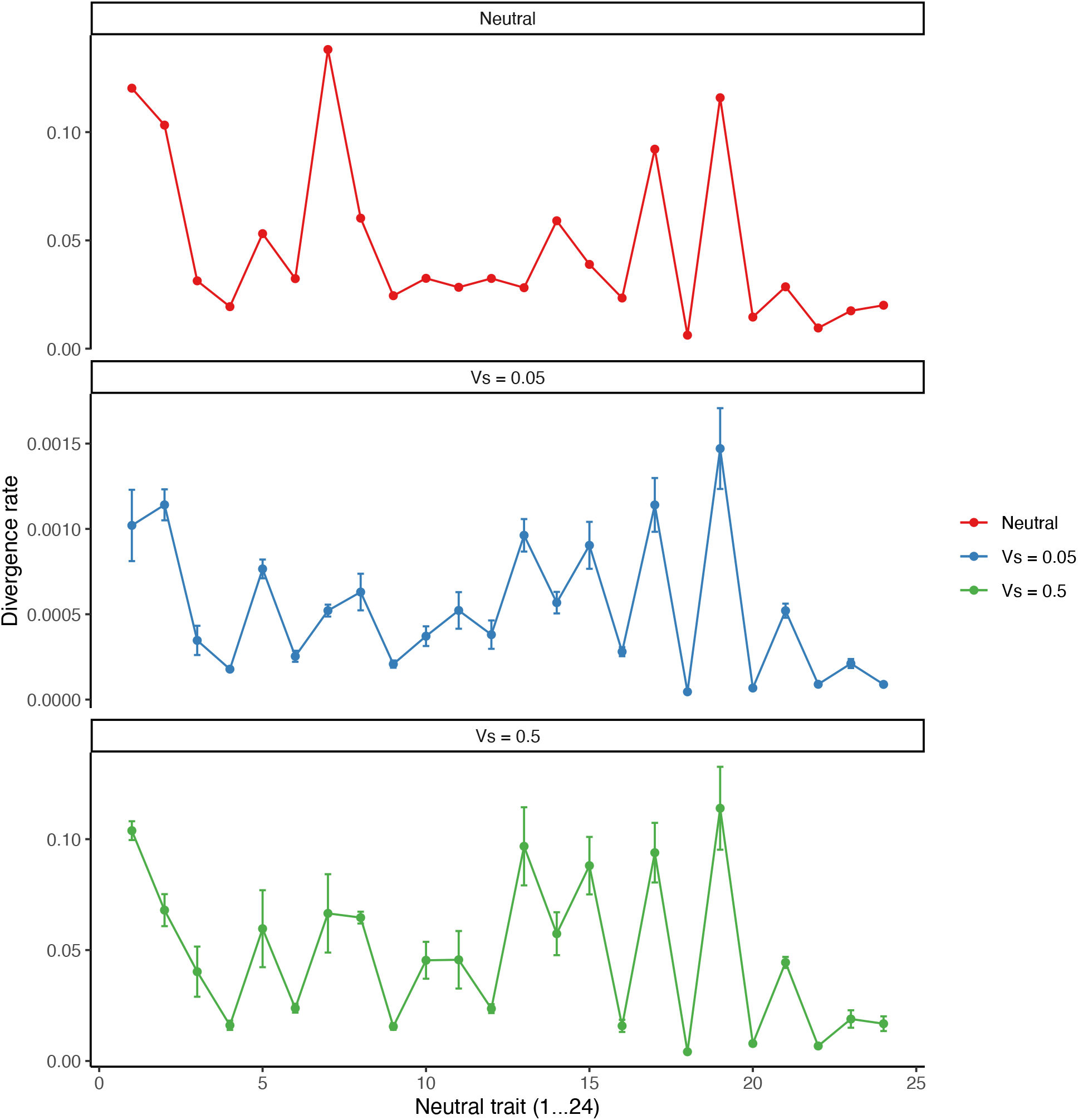
Divergence rate for 24 non-selected wing shape traits under single-axis selection. Each point represents the divergence rate (increase in among-population variance per generation) for one wing shape trait. Under strong stabilizing selection on wing size (*V*_*s*_ = 0.05, blue), divergence rates are substantially reduced compared to weak selection (*V*_*s*_ = 0.5, green) and the completely neutral scenario (red, where both wing size and shape are neutral). This indicates that selection on a single primary trait can constrain the evolution of non-selected traits. Simulations used *σ* = 0.0001, *N* = 500, and 20 replicate populations evolved for 20,000 generations. Three independent simulation replicates were used for each *V*_*s*_ condition (*V*_*s*_ = 0.05 and *V*_*s*_ = 0.5).

**Figure 4:**
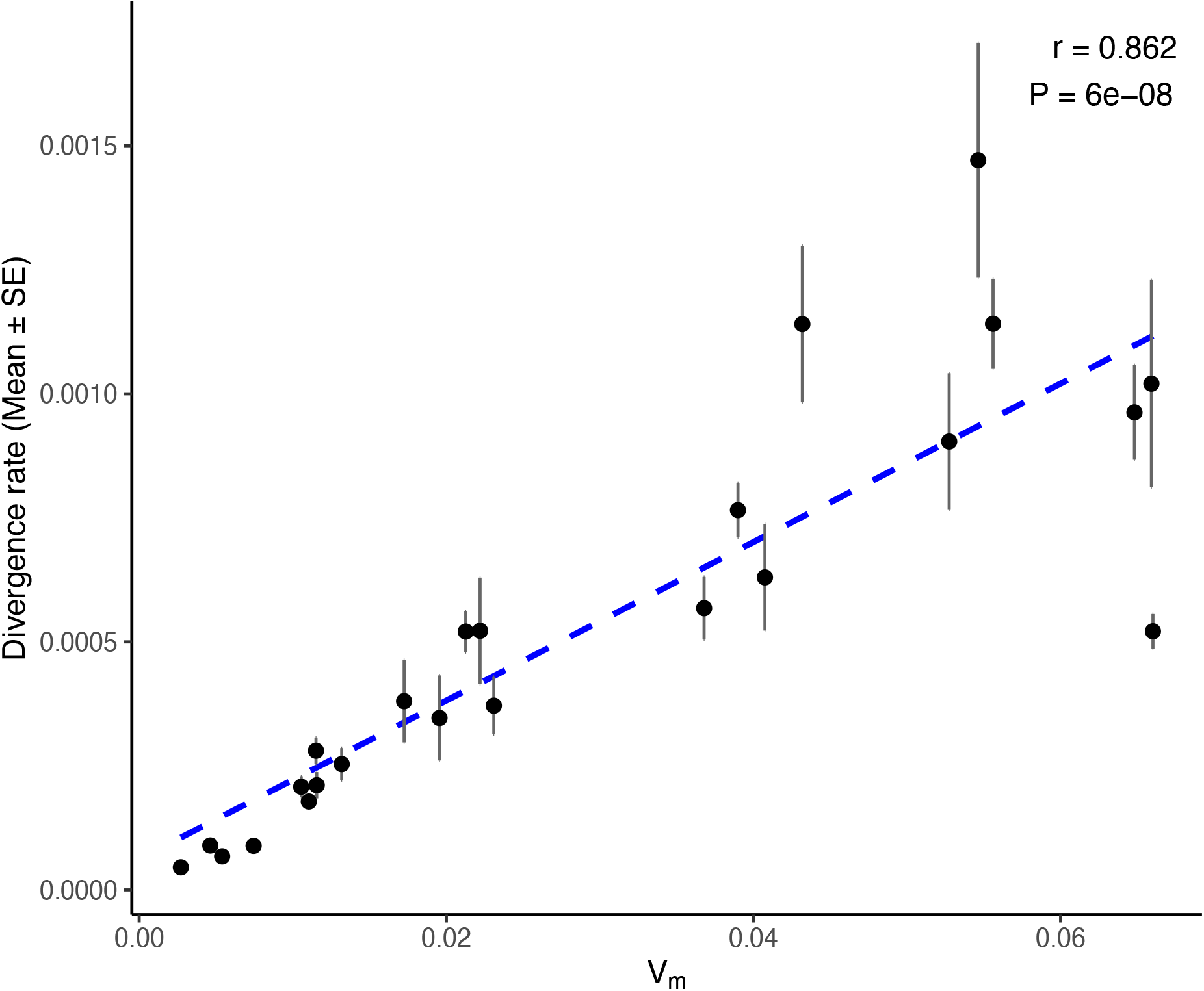
Divergence rate for non-selected traits scales with mutational variance under single-axis selection. Each point represents one of the 24 shape traits. The positive relationship between mutational variance (*V*_*M*_) and divergence rate shows that traits with higher mutational input diverge faster, even when most traits are not under direct selection. This scaling underlies the alignment between *M, G*, and *R*. Simulations used *V*_*s*_ = 0.05 on wing size, *σ* = 0.0001, *N* = 500, and 20 replicate populations. Three independent simulation replicates were used.

### Single-Axis Selection and Correlational Selection Leave Distinct Imprints on *G*

The single-axis selection model and the correlational selection hypothesis make distinct predictions about the eigenvalue structure of *G* relative to *M* . If correlational selection is the primary force shaping genetic covariances—favoring specific trait combinations that mirror the mutational architecture—then the equilibrium *G* should preserve the eigenvalue structure of *M*, maintaining similar eccentricity (i.e., similar ratios among eigenvalues). Conversely, under single-axis selection, where only a single primary trait is under selection, variance along the leading eigenvector of *M* (which is size-dominated) should be preferentially eroded, while variance in orthogonal directions remains largely unconstrained. This predicts that *G* should be less eccentric than *M* (here, size is included in the matrices).

Ref. ^15^ found that mutational correlations among fly wing traits are consistently stronger than the corresponding additive genetic correlations. Furthermore, mutational variance is highly concentrated along a single multivariate axis (high eccentricity), while genetic variance is more evenly distributed across axes (lower eccentricity). Reanalysis of *G* and *M* from ref. ^9^ reveals a similar pattern: the ratio of the first eigenvalue is substantially larger for *M* than for *G* (Fig. S5 and Appendix B). These independent datasets converge on a consistent finding: selection reduces, rather than reinforces, the eccentricity imposed by mutational input.

To formally test these competing predictions, I conducted simulations comparing equilibrium *G* under two scenarios (using *M* obtained from ref. ^15^): (1) the single-axis selection model, where only the primary trait is under stabilizing selection; and (2) correlational selection, where the selection matrix *S* is proportional to *M* ^−1^, imposing multivariate stabilizing selection that favors the trait combinations encoded by mutational pleiotropy. Two quantitative diagnostics were applied to the simulated equilibrium *G* matrices (5 replicates per scenario) to formally discriminate the two hypotheses.

The results (Fig. 6 and Fig. S12) show that the eigenvalue spectrum of equilibrium *G* under single-axis selection is visibly less eccentric than that of *M* : the proportion of total genetic variance explained by the leading eigenvalue is reduced under single-axis selection, with the residual variance redistributed among the rest of the eigenvectors; in contrast, correlational selection preserves the eigenvalue proportions close to those of *M* . This flattening is also captured by the effective dimensionality of *G* (*n*_eff_; ref. ^16,17^, See Material and Methods): under single-axis selection, *n*_eff_ = 2.17 ± 0.35 at *V*_*s*_ = 1 and 2.09 ± 0.15 at *V*_*s*_ = 5 (mean ± s.d. across replicates), similar to empirical *G*, all exceeding the *M* baseline (*n*_eff_ = 1.80, computed directly from the mutational covariance matrix), whereas correlational selection leaves *n*_eff_ essentially unchanged (1.79 ± 0.19 at *V*_*s*_ = 1; 1.88 ± 0.26 at *V*_*s*_ = 5). These diagnostics collectively suggest that selection on a single trait reshapes the eigenvalue structure of *G* in a manner qualitatively distinct from multivariate correlational selection. Notably, *n*_eff_ empirical *G* exceeds the simulated single-axis values at all selection strengths simulated (Fig. S13), indicating that additional factors— such as environmental variance, selection on multiple targets, or more complex pleiotropic architectures— contribute to the dimensionality of empirical *G* beyond what the idealized single-trait model predicts.

These findings should be interpreted cautiously, given that empirically estimated *M* matrices may carry greater sampling uncertainty due to lower effective sample sizes ^15^, and the two scenarios tested here only represent idealized extremes of a continuum that may include both processes operating simultaneously.

## Discussion

The strong alignment between mutational variation (*M*), genetic variation (*G*), and species divergence (*R*) in fly wing shape has puzzled evolutionary biologists for nearly a decade. Here, I show that a minimal single-axis selection model— in which natural selection targets only one trait while all other traits evolve as correlated byproducts through within-module pleiotropy— is sufficient to reproduce these empirical patterns without invoking hidden constraints, correlational selection, or complex adaptive landscapes. The model predicts that the directly selected trait should exhibit the lowest *G/M* ratio, relative to other non-selected traits. Reanalysis of empirical *M* and *G* reveals that wing size may indeed be the primary target of selection.

### Evidence for Wing Size as the Primary Selection Target

The single-axis selection model requires that wing size experiences stronger direct selection than wing shape. There is considerable evidence supporting this assumption. Wing size is a critical determinant of aerodynamic performance, directly influencing lift generation and power requirements for flight. Research has documented selection on wing size through multiple fitness components: mating success ^18–20^, migration survival ^21,22^, thermoregulation ^23^, and fecundity ^24–27^. In sepsid flies specifically, a comparative study of sexual selection across seven species found that larger males consistently had higher mating success, and that sexual selection on wing and fore femur shape was closely aligned with static allometry^20^. This alignment is consistent with the hypothesis that apparent selection on shape is a byproduct of selection on overall body size. However, allometry-corrected shape selection often remained significant ^20^, indicating that some selection on shape persists independently of size. Because the size-related mating differentials were not corrected for correlated shape variation, the converse interpretation—that selection targets shape and produces apparent selection on size through allometry—cannot be fully excluded. Note that optimal wing size likely depends on body mass. Therefore, at the whole-organism level, selection is never ‘single-axis’ and on size alone: when contrasting “single-axis selection” with “correlational selection” in this study, only the evolutionary dynamics within the wing module are considered, encompassing the 25 traits available in the dataset.

Direct evidence for selection on the relative positions of wing vein intersections (i.e., shape independent of size) is scarce. While vein positioning may influence wing flexibility and aerodynamic properties ^28^, the functional consequences of natural shape variation appear subtle compared to size effects. In odonates, wing shape does show significant genetic differentiation across latitudinal gradients, with genetic population differences having two to four times the impact of phenotypic plasticity on shape variation^29^. However, even in these systems, wing shape variation may be tightly linked to life-history traits through developmental constraints and pleiotropy^29^; body size, which varies clinally with developmental time constraints^26^, can scale allometrically with wing shape. These findings suggest that size-shape developmental integration provides a general mechanism by which selection on size can indirectly constrain shape evolution, although direct selection on shape cannot be entirely excluded.

In the empirical *M* matrix used for the simulation, wing size carries roughly two to three orders of magnitude more mutational variance than the average shape trait, raising the question of whether the alignment result depends on selecting this dominant axis. Repeating the simulation with selection on a low-variance shape trait (trait 1) instead of size confirmed that the alignment is not contingent on selecting the highest-variance axis (Appendix C, Fig. S8). However, empirical data shows that wing size has the lowest *G/M* ratio of all 25 traits (Fig. 1). As established earlier, this trait-specific depletion profile may serve as a diagnostic test for size being the primary selection target, rather than a shape trait (Fig. 5 and Fig. S9). This robustness test verifies that the apparent selection mechanism is general, while providing support for size as the primary selection target.

**Figure 5:**
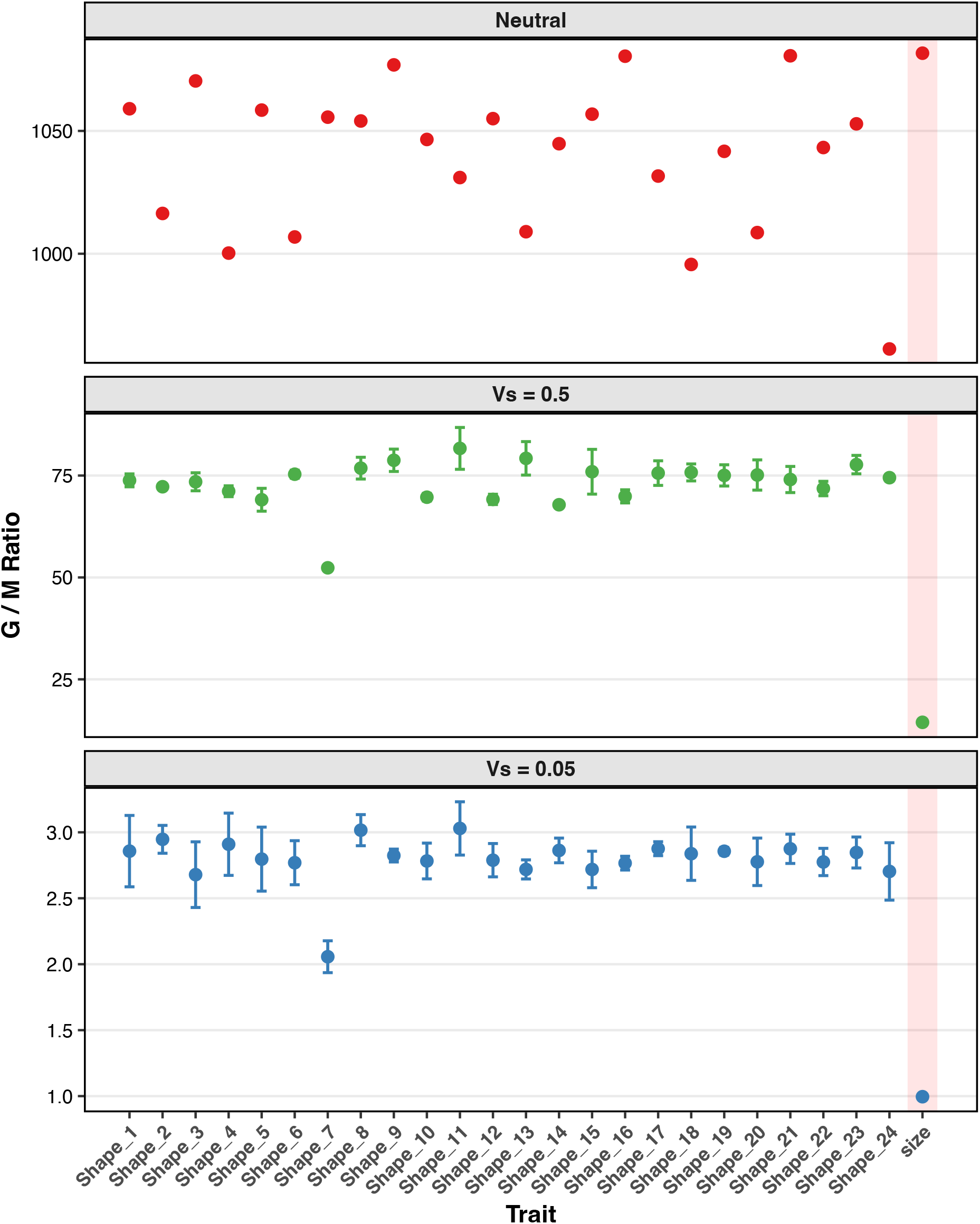
Ratio of genetic to mutational variance (*G/M*) across traits under varying evolutionary scenarios. Points represent the mean *G/M* ratio across replicates for each trait under pure neutral drift, weak stabilizing selection on the primary trait (*V*_*s*_ = 0.5), and strong stabilizing selection on the primary trait (*V*_*s*_ = 0.05). Error bars indicate± 1 standard error of the mean across independent simulation replicates (*n* = 3 for selection scenarios). The neutral panel reflects a single baseline simulation (*n* = 1) and thus lacks error bars. The focal “size” trait is highlighted with a red shaded background to facilitate direct visual comparison of its variance ratio relative to other traits. Note that the y-axis scales vary independently across panels to accommodate differing magnitudes of *G/M* ratios between scenarios.

**Figure 6:**
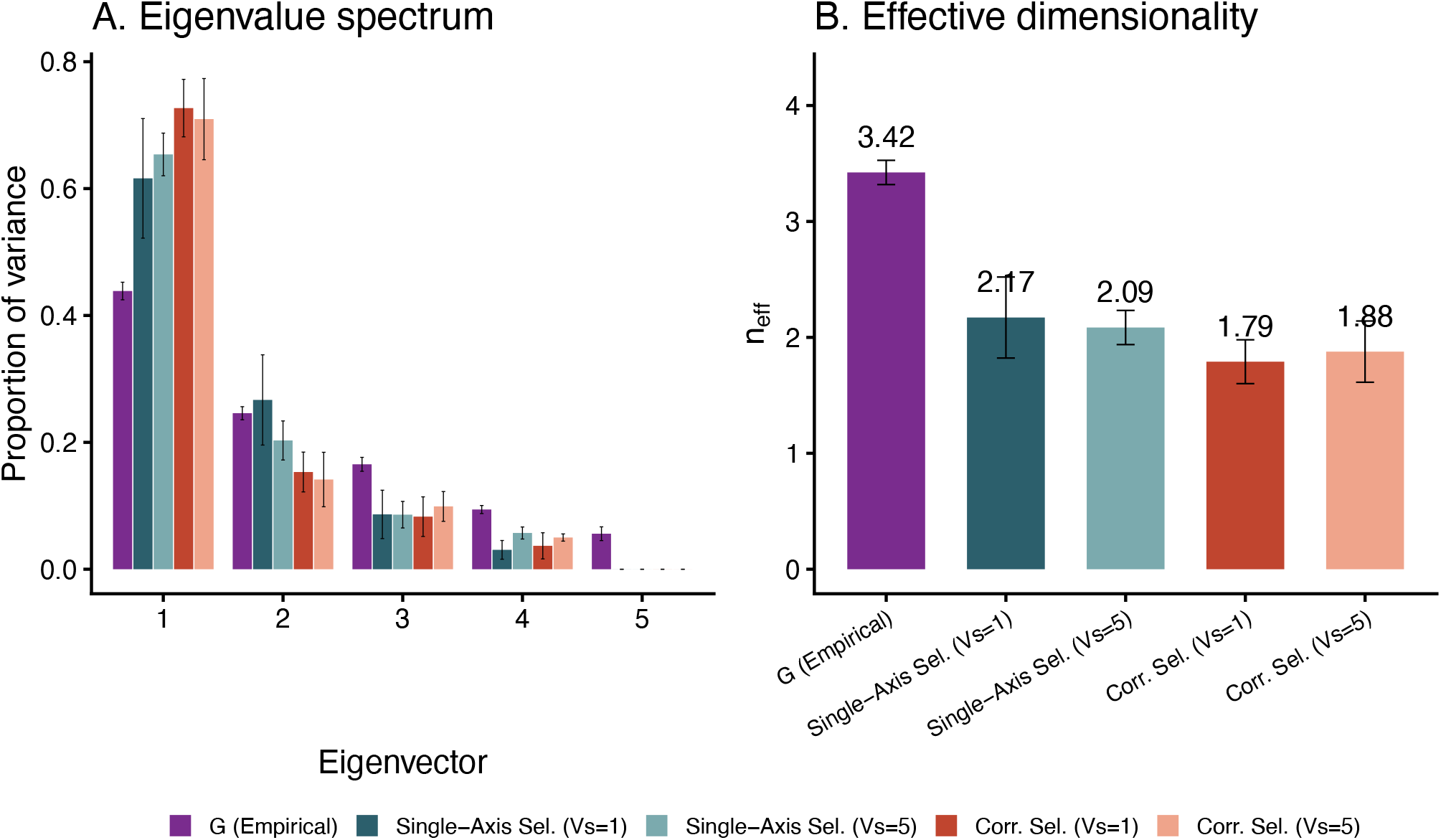
Eccentricity diagnostics for simulated and empirical *G* matrices. Two diagnostics applied to the empirical *G* from ref. ^15^ and equilibrium *G* under single-axis selection and correlational selection at two selection strengths (*V*_*s*_ = 1 and *V*_*s*_ = 5; 5 replicates each, *N* = 500). (A) Eigenvalue spectrum showing the proportion of total genetic variance explained by each eigenvector. (B) Effective dimensionality (*n*_eff_), measuring how evenly variance is distributed across eigenvectors. Error bars for simulated *G* show ±1 s.d. across replicates; error bars for empirical *G* represent estimation uncertainty (parametric bootstrap from REML confidence intervals reported in ^15^).

### Stability of the Mutational Variance-Covariance Matrix

A key assumption of the single-axis selection model is that the structure of *M* remains stable over macroevolutionary time. Several lines of evidence support this assumption. First, genetic variance-covariance matrices (*G*) appear similar between Sepsidae and *Drosophila melanogaster*, which diverged approximately 64 million years ago ^10,30^. Common subspace analyses comparing *G* matrices across these taxa reveal substantial alignment regardless of which species is used as the reference (Figs. S10, S11). However, because *G* is determined by both mutational input and selection, observing that *G* is conserved does not strictly imply that *M* is conserved. If selection is strong and conserved, it could constrain *G* to remain stable even if the underlying mutational architecture fluctuates. Inferring that *M* is conserved from a conserved *G* requires the assumption that selection is not completely overriding the mutational variance. Second, the highly polygenic genetic architecture of wing shape traits (Appendix E) may enhance the stability of *M* and *G* by buffering against the effects of individual mutations ^31,32^. Nevertheless, direct measurement of *M* across divergent lineages remains an important goal for future research.

### Ultimate versus Proximate Causes

It is important to distinguish between proximate and ultimate explanations for the *M* -*G*-*R* alignment. The single-axis selection model provides a proximate mechanism: given a stable *M* matrix and selection on a single trait, alignment emerges as a natural consequence. Similarly, the multidimensional fluctuating-selection (MFS) model^3^operates as an alternative proximate mechanism: assuming a pre-existing mutational and genetic architecture, it demonstrates how rapid fluctuations in optima produce macroevolutionary divergence (*R*) that aligns with *G*. However, the ultimate origin of the mutational architecture itself remains an open question. Mutation is evolvable ^4,33^, and the structure of *M* may itself be shaped by selection over deep evolutionary time.

Several ultimate-level hypotheses offer explanations for the origin of *M* . The “moving corridor” hypothesis proposes that the mutational architecture itself evolves to align with the predominant direction of optimum movement: lineages whose *M* channels variation along the historical axis of environmental change adapt more readily, so over deep time *M* comes to mirror the typical trajectory of shifting optima. Correlational selection models propose that persistent selection on trait combinations (e.g., allometric relationships) shapes *M* and *G* simultaneously.

### Generality beyond Fly Wings

The results presented here focus on the *Drosophila* wing, but the features of the present model and the underlying mechanism should apply to trait systems exhibiting stable within-module pleiotropy. For example, macroevolutionary divergence predominantly tracking a single size or allometric line of least resistance is a widespread phenomenon, documented in ruminant skulls^34^, amphibian locomotion ^35^, and morphological differentiation in *Pristurus* geckos ^36^. Yet, in systems where morphological features are governed by largely independent developmental modules, like absolute scalar size and geometric shape in Darwin’s finch beaks ^37^, natural selection can act on these axes independently.

The key prediction of this framework is thus not that selection always targets a single axis like size, but that within-module pleiotropy with low effective dimensional selection should exhibit the hallmarks identified here: divergence of non-selected traits depending on the axis under selection and depletion of standing genetic variance concentrated primarily on the axis under selection. Testing these predictions across diverse morphological modules would provide a powerful assessment of how broadly the correlated-response mechanism contributes to observed alignments between micro- and macroevolution^8^.

## Materials and Methods

### Model Overview

The single-axis selection model, proposed here, posits that natural selection targets only one primary trait, while remaining traits evolve as neutral byproducts through their mutational covariance with the selected trait. The instantaneous selection gradient *β* implied by the Gaussian fitness function is therefore:

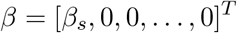

where *β*_*s*_ reflects the strength of selection on the primary trait, determined by the distance between the population mean and the current optimum. The population divergence in the primary trait arises from lineage-specific moving optima. In this reduced wing-only model, the dependence of optimal wing size on body mass and environments is represented as a lineage-specific moving optimum for wing size. Under this formulation, wing shape traits are effectively neutral, but experience indirect evolutionary change through their mutational covariance with the selected trait.

### Individual-Based Simulations

I assessed the single-axis selection model using individual-based simulations. I simulated populations of *N* = 500 hermaphroditic, sexually reproducing individuals with non-overlapping generations. Phenotypes were determined by *n* = 50 unlinked diploid loci, with alleles assumed to be fully pleiotropic (i.e., affecting all phenotypic traits). The life cycle proceeded in three discrete steps:

1. **Phenotype and fitness assignment**. Each individual’s phenotypic value for every trait was obtained by summing allelic breeding values across all loci, assuming strict additivity (no dominance or epistasis at the phenotypic level, though epistasis may arise at the fitness level). Fitness was then assigned according to a Gaussian stabilizing selection function applied solely to the primary trait:

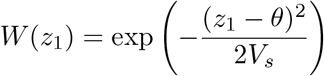

where *z*_1_ is the individual’s primary trait value, *θ* is the current phenotypic optimum, and *V*_*s*_ controls the width of the fitness peak (smaller *V*_*s*_ imposes stronger selection).
2. **Selection and mating**. Parents were sampled with replacement in proportion to their fitness, and two distinct individuals were paired to produce each offspring (selfing was excluded). This fitness-weighted sampling was repeated *N* times per generation to maintain constant population size.
3. **Recombination and mutation**. Offspring genotypes were assembled by independently drawing one allele per locus from each parent, corresponding to free recombination among all loci. Mutations occurred at a rate of *µ* = 0.01 per allele per generation (genomic mutation rate *U* = 1 per individual per generation). Pleiotropic mutational effects were drawn from a multivariate normal distribution with variance-covariance matrix **M**, estimated empirically under homozygous conditions by ref. ^9^.

Each simulation began with a burn-in period of 10*N* generations, with the optimum set to zero throughout. Following burn-in, 20 replicate populations were evolved for *T* = 20000 generations under stabilizing selection on lineage-specific moving optima for the primary trait, where each replicate was assigned an optimum moving rate sampled from a normal distribution with mean *µ* = 0 and standard deviation *σ* = 0.0001. Selection strength was varied across simulations by setting *V*_*s*_ = 0.05 (strong selection) or *V*_*s*_ = 0.5 (weak selection).

The within-population genetic variance-covariance matrix **G** was computed as the phenotypic covariance matrix within each replicate population at the final generation and then averaged across all 20 replicates. The among-population divergence matrix **R** was computed as the covariance matrix of the 20 replicate-population trait means.

Following ref. ^9^, centroid size enters both the empirical **M** and **G** matrices as its natural logarithm (ln CS). The *G/M* ratio is intrinsically dimensionless, making it invariant to measurement scale. It is also robust to the log-transformation: by the first-order delta method, Var 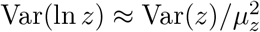 and because this scaling factor applies equally to both variances, *G*_ln *z*_*/M*_ln *z*_ *≈ G*_*z*_*/M*_*z*_.

### Comparing Variance-Covariance Matrices

To quantify the alignment between mutational variance (**M**), genetic variance (**G**), and evolutionary divergence (**R**), I employed the common subspace analysis of ref. ^14^. This approach compares matrices along a shared set of orthogonal axes defined by the eigenvectors of **M**.

Specifically, I decomposed the homozygous mutational variance-covariance matrix (**M**_*hom*_) estimated in *D. melanogaster* into its eigenvector matrix **K**. For each variance-covariance matrix of interest (**X** ∈ {**G, R**}), I calculated the variance along each eigenvector as the diagonal entries of **K**^T^**XK**. I then regressed log_10_-transformed variances in **R** or **G** against log_10_-transformed variances in **M** using ordinary least squares (OLS), obtaining regression slopes and coefficients of determination (*R*^2^). To avoid comparisons along dimensions with negligible variance, analyses were restricted to the first 18 eigenvectors of **M**.

### Eccentricity Diagnostics

To discriminate between the single-axis selection and correlational selection hypotheses, I applied two quantitative diagnostics to the eigenvalue structure of the equilibrium genetic variance-covariance matrix *G* relative to the mutational variance-covariance matrix *M* .

The first diagnostic is the eigenvalue spectrum, which compares the proportion of total genetic variance captured by each successive eigenvector of *G*. Under single-axis selection, selection on the primary trait preferentially erodes variance along the leading eigenvector of *M* (which is dominated by size), redistributing variance toward trailing eigenvectors and flattening the spectrum. Under correlational selection, the eigenvalue proportions of *G* should remain similar to those of *M* .

The second diagnostic is the effective dimensionality, which is defined as participation ratio^16,17^:

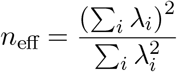

where *λ*_*i*_ are the eigenvalues of *G*. This index measures how evenly variance is distributed across eigenvectors: *n*_eff_ = 1 when all variance is concentrated along a single axis, and *n*_eff_ = *n* when variance is uniformly distributed across all *n* eigenvectors. This definition of effective dimensionality is different from the one used in ref. ^38^. Yet, all the results for *n*_eff_ are qualitatively similar regardless which definition is used. Single-axis selection is predicted to increase *n*_eff_ above the *M* baseline by reducing the dominance of the leading eigenvalue.

For each scenario, equilibrium *G* matrices were obtained from individual-based simulations using the empirical *M* matrix of ref. ^15^ (5 traits: centroid size plus 4 shape variables). The empirical *M* was forced to be positive semi-definite prior to use. Simulations were run with *N* = 500 individuals for 5 independent replicates per scenario. Because the ref. ^15^ traits are on a different measurement scale (due to different traits and scaling methods during analyses) from those of ref. ^9^, the stabilizing selection width *V*_*s*_ was adjusted. For the empirical *G* matrix (ref. ^15^), estimation uncertainty was quantified by parametric bootstrap: 200 replicate matrices were generated by sampling each element from a normal distribution centered on the point estimate with standard deviation equal to the reported standard error, followed by symmetrization and projection onto the nearest positive semi-definite matrix.

## Acknowledgements

I thank two anonymous reviewers for their careful readings and constructive comments that substantially improved the manuscript. I thank Patrick Rohner for sharing the data used in Appendix D and for valuable discussions. I also thank David Houle, Daohan Jiang, Joel McGlothlin and Michael Whitlock for helpful discussions.

## References

[1] Rolland, J. et al. Conceptual and empirical bridges between micro-and macroevolution. Nature Ecology & Evolution 7, 1181–1193 (2023).

[2] Pavlicev, M. Cracking open the blackbox of genotype-phenotype map: Crossing the explanatory gap between micro-and macroevolution. Evolution (2025).

[3] Holstad, A. et al. Evolvability predicts macroevolution under fluctuating selection. Science 384, 688–693 (2024).

[4] Cai, H., Melo, D. & Des Marais, D. L. Disentangling variational bias: the roles of development, mutation, and selection. Trends in Genetics 41, 23–32 (2025). URL https://www.sciencedirect.com/science/article/pii/S0168952524002300.

[5] Saito, K., Tsuboi, M. & Takahashi, Y. Conserved wing shape variation across biological scales unveils dialectical relationships between micro-and macroevolution. Communications Biology 8, 990 (2025).

[6] Melo, D., Porto, A., Cheverud, J. M. & Marroig, G. Modularity: genes, development, and evolution. Annual review of ecology, evolution, and systematics 47, 463–486 (2016).

[7] Tsuboi, M. et al. The paradox of predictability provides a bridge between micro-and macroevolution. Journal of evolutionary biology 37, 1413–1432 (2024).

[8] Schluter, D. Variable success in linking micro-and macroevolution. Evolutionary Journal of the Linnean Society 3, kzae016 (2024).

[9] Houle, D., Bolstad, G. H., van der Linde, K. & Hansen, T. F. Mutation predicts 40 million years of fly wing evolution. Nature 548, 447–450 (2017).

[10] Rohner, P. T. & Berger, D. Macroevolution along developmental lines of least resistance in fly wings. Nature Ecology & Evolution 1–13 (2025).

[11] Pennell, M. & Jiang, D. The macroevolutionary adaptive landscape: more than a metaphor? (2024).

[12] Arnold, S. J. Evolutionary quantitative genetics (Oxford University Press, 2023).

[13] Houle, D., Bolstad, G. H. & Hansen, T. F. Fly wing evolutionary rate is a near-isometric function of mutational variation. bioRxiv 2020–08 (2020).

[14] Jiang, D. & Zhang, J. Fly wing evolution explained by a neutral model with mutational pleiotropy. Evolution 74, 2158–2167 (2020).

[15] Dugand, R. J., Aguirre, J. D., Hine, E., Blows, M. W. & McGuigan, K. The contribution of mutation and selection to multivariate quantitative genetic variance in an outbred population of drosophila serrata. Proceedings of the National Academy of Sciences 118, e2026217118 (2021).

[16] Wang, Z. et al. The geometry and dimensionality of brain-wide activity. Elife 14, RP100666 (2025).

[17] Gao, P. et al. A theory of multineuronal dimensionality, dynamics and measurement. BioRxiv 214262 (2017).

[18] Yenisetti, S. C. & Hegde, S. N. Size-related mating and reproductive success in a drosophilid: Phorticella striata. ZOOLOGICAL STUDIES-TAIPEI-42, 203–210 (2003).

[19] Kotyk, M. & Varadínová, Z. Wing reduction influences male mating success but not female fitness in cockroaches. Scientific reports 7, 2367 (2017).

[20] Blanckenhorn, W. U. et al. Comparative sexual selection in field and laboratory in a guild of sepsid dung flies. Animal Behaviour 175, 219–230 (2021).

[21] Altizer, S. & Davis, A. K. Populations of monarch butterflies with different migratory behaviors show divergence in wing morphology. Evolution 64, 1018–1028 (2010).

[22] Dockx, C. Directional and stabilizing selection on wing size and shape in migrant and resident monarch butterflies, danaus plexippus (l.), in cuba. Biological Journal of the Linnean Society 92, 605–616 (2007).

[23] Berwaerts, K., Van Dyck, H., Vints, E. & Matthysen, E. Effect of manipulated wing characteristics and basking posture on thermal properties of the butterfly pararge aegeria (l.). Journal of Zoology 255, 261–267 (2001).

[24] Chang, H., Guo, X., Guo, S., Yang, N. & Huang, Y. Trade-off between flight capability and reproduction in acridoidea (insecta: Orthoptera). Ecology and Evolution 11, 16849–16861 (2021).

[25] McCulloch, G. A. et al. Dispersal-fecundity trade-offs in wild insect populations. Journal of evolutionary biology 38, 430–436 (2025).

[26] Outomuro, D., Golab, M. J., Johansson, F. & Sniegula, S. Body and wing size, but not wing shape, vary along a large-scale latitudinal gradient in a damselfly. Scientific Reports 11, 18642 (2021).

[27] Yang, J., Yang, C., Lin, H.-w., Lees, A. C. & Tobias, J. A. Elevational constraints on flight efficiency shape global gradients in avian wing morphology. Current Biology 35, 1890–1900 (2025).

[28] Combes, S. A. & Daniel, T. L. Flexural stiffness in insect wings i. scaling and the influence of wing venation. Journal of experimental biology 206, 2979–2987 (2003).

[29] Johansson, F. et al. Mixed support for an alignment between phenotypic plasticity and genetic differentiation in damselfly wing shape. Journal of Evolutionary Biology 36, 368–380 (2023).

[30] Wiegmann, B. M. et al. Episodic radiations in the fly tree of life. Proceedings of the National Academy of Sciences 108, 5690–5695 (2011).

[31] Barton, N. H. & Turelli, M. Evolutionary quantitative genetics: how little do we know. Annual review of genetics 23, 337–370 (1989).

[32] Cai, H., Geiler-Samerotte, K. & Des Marais, D. L. Dissecting genetic correlation through recombinant perturbations: the role of developmental bias. bioRxiv 2023–05 (2023).

[33] Jones, A. G., Bürger, R. & Arnold, S. J. Epistasis and natural selection shape the mutational architecture of complex traits. Nature communications 5, 3709 (2014).

[34] Rhoda, D. P., Haber, A. & Angielczyk, K. D. Diversification of the ruminant skull along an evolutionary line of least resistance. Science Advances 9, eade8929 (2023).

[35] Simon, M. N., Courtois, E. A., Herrel, A. & Moen, D. S. Macroevolutionary divergence along allometric lines of least resistance in frog hindlimb traits and its effect on locomotor evolution. The American Naturalist 205, 637–655 (2025).

[36] Tejero-Cicuéndez, H. et al. Evolution along allometric lines of least resistance: morphological differentiation in pristurus geckos. Evolution 77, 2547–2560 (2023).

[37] Grant, P. R. & Grant, B. R. 40 years of evolution: Darwin’s finches on daphne major island (2014).

[38] Kirkpatrick, M. Patterns of quantitative genetic variation in multiple dimensions. Genetica 136, 271–284 (2009).

